# Top-down specific preparatory activations for Selective Attention and Perceptual Expectations

**DOI:** 10.1101/2022.09.13.507583

**Authors:** José M. G. Peñalver, David López-García, Carlos González-García, Blanca Aguado-López, Juan M. Górriz, María Ruz

**Affiliations:** Mind, Brain and Behavior Research Center (CIMCYC), University of Granada, Granada 18071, Spain; Data Science & Computational Intelligence Institute, Universidad de Granada, CP 18071; Cambridge Neuroscience, University of Cambridge, CB2 0SZ, UK

**Keywords:** Selective Attention, Expectations, Preparation, EEG, MVPA, RSA

## Abstract

**Summary:** Proactive cognition brain models are mainstream nowadays. Within these, preparation is understood as an endogenous, top-down function that takes place prior to the actual perception of a stimulus and improves subsequent behavior. Neuroimaging has shown the existence of such preparatory activity separately in different cognitive domains, however no research to date has sought to uncover their potential similarities and differences. Two of these, often confounded in the literature, are Selective Attention (information relevance) and Perceptual Expectation (information probability). We used EEG to characterize the mechanisms that pre-activate specific contents in Attention and Expectation. In different blocks, participants were cued to the *relevance* or to the *probability* of target categories, faces vs. names, in a gender discrimination task. Multivariate Pattern (MVPA) and Representational Similarity Analyses (RSA) during the preparation window showed that both manipulations led to a significant, ramping-up prediction of the relevant or expected target category. However, classifiers trained on data from one condition did not generalize to the other, indicating the existence of unique anticipatory neural patterns. In addition, a Canonical Template Tracking procedure showed that there was stronger anticipatory perceptual reinstatement for relevance than for expectation blocks. Overall, results indicate that preparation during attention and expectation acts through distinguishable neural mechanisms. These findings have important implications for current models of brain functioning, as they are a first step towards characterizing and dissociating the neural mechanisms involved in top-down anticipatory processing.

## Introduction

For decades, research in Cognitive Psychology has studied behavior while manipulating external factors, which led to theoretical models that framed cognition mostly from a reactive point of view. Recent years have witnessed a renaissance of proactive cognition, where endogenous top-down mechanisms play a core role in brain functioning. Within this framework, preparation is conceptualized as an endogenous function that takes place in anticipation of incoming inputs or demands and improves subsequent behavior (Battistoni et al., 2017; González-García et al., 2016). Neural preparatory activity has been shown for a plethora of processes, including attention (Battistoni et al., 2017; Kastner et al., 1999; Nobre & Serences, 2018), expectation (Aranda et al., 2010; Kok et al., 2017), working memory (Koshino et al., 2015; van Driel et al., 2017), or cognitive control (Baines et al., 2011; González-García et al., 2016; Hebart & Baker, 2018). Similarly, influential models have proposed different ways in which anticipatory patterns interact with stimulus inputs to guide perception. Examples of this are the Predictive Coding (Auksztulewicz & Friston, 2016a) or the Biased Competition frameworks (Desimone & Duncan, 1995). However, currently these models are silent as to how different preparatory phenomena relate to each other, and whether they reflect common or diverging underlying top-down mechanisms.

Selective Attention and Perceptual Expectation are complex functions that involve top-down and bottom-up elements. Attention refers to the selection of *relevant* information based on specific goals (Nobre & Serences, 2018), while expectation involves predictions based on prior *probability* (Schröger et al., 2015). Studies of selective attention have manipulated the relevance of information using cues that indicate the stimulus or dimensions to respond while ignoring others (Battistoni et al., 2017; Hong et al., 2017; Nobre & Serences, 2018; Stokes et al., 2009). Expectation has primarily (but not only, see e.g. Summerfield & De Lange, 2014) been manipulated with cues that inform about the most probable stimulus (de Lange et al., 2013; Kok et al., 2017; Wyart et al., 2012a). Previous research focusing on the effect of Attention and Expectations on target processing has shown diverging (brain activity) and overlapping (behavioral) results. Although both lead to behavioral improvements (Ho et al., 2012; Stein & Peelen, 2015), several studies reveal that they can be, at least partially differentiable (Rungratsameetaweemana & Serences, 2019; Summerfield & Egner, 2009). Neuroimaging studies so far have found differences in contexts of relevance and probability (see Summerfield & Egner, 2009, 2016 for a summary), including activity increases for selected target stimuli and decreases for expected ones. Another fruitful field of research has focused on how they interact, sowing in some cases how selective attention can modulate the effect of expectations (Alilović et al., 2019; Auksztulewicz et al., 2017; Jiang et al., 2013; Kok et al., 2012) and a lack of interactions in others (e.g. Ekman et al., 2017; Yon et al., 2018).

Studies that have tried to overcome frequent confounds between attention and expectation (e.g. Posner, 1980; Schröger, 1996)) have shown separate roles of relevance and probability during target processing (Auksztulewicz et al., 2017; Gordon et al., 2019; Simon et al., 2019; Wyart et al., 2012b; Zuanazzi & Noppeney, 2019). On the other hand, research focused on preparatory activity of either attention or expectation has provided seemingly overlapping results. Cues indicating relevance in selective attention (Battistoni et al., 2017; Nobre & Serences, 2018) preactivate relevant regions of space processing (Giesbrecht et al., 2006), specific shape patterns in visual cortex (Stokes et al., 2009), patterns in category (Esterman & Yantis, 2010; González-García et al., 2018) and object-selective perceptual regions (Peelen & Kastner, 2011; Soon et al., 2013). Similarly, cues providing probabilistic information lead to the preactivation of specific templates of oriented gabors (Kok et al., 2017), direction (Ekman et al., 2017), motor patterns (de Lange et al., 2013) or abstract shapes (Hindy et al., 2016). However, all these previous investigations are agnostic regarding the potential similarities or differences in such top-down preparation across relevance and probability anticipation. Unraveling the differences in how anticipatory activity in different contexts reflects the upcoming information is a necessary step to understand the differences between Attention and Expectation, and is also essential for theoretical models that explain the neural basis of these two phenomena (Auksztulewicz et al., 2018; de Lange et al., 2018; Desimone & Duncan, 1995).

In our study, we employed different multivariate pattern analyses of EEG data to directly compare the representational structure of anticipated contents for top-down preparation in contexts capitalizing on Selective Attention (relevance) or Expectation (probability). That is, we examined whether the coding of anticipated stimulus content differs on the basis of the type of anticipation. To do so, we embedded a sex/gender discrimination in a cue-target paradigm. Here, depending on the block, cues provided information about the upcoming relevance or probability (Egner et al., 2010; Wyart et al., 2012b) of face or name stimulus categories. In addition, we ran an independent localizer to study similarities between preparation and perception across contexts. Our first goal was to study the anticipatory mechanisms during preparation by means of time-resolved representational similarity analysis (RSA, Kriegeskorte, 2008). Then, we used multivariate pattern analysis (MVPA, Grootswagers et al., 2017) to examine whether anticipated stimulus categories are represented with differential fidelity during selected compared to probable targets. Next, we used a cross-classification approach to directly contrast the patterns of activity underlying the representation of relevant vs. probable stimuli. Taking into consideration the striking differences observed during target processing in these contexts (Jiang et al., 2013; Kok et al., 2012; Summerfield & Egner, 2009; Wyart et al., 2012a), our hypothesis was that the neural coding that leads to such different consequences should be dissociable from an early processing stage. To further understand the proposed differences, we leveraged a Canonical Template Tracking approach (González-García et al., 2021a; Palenciano et al., 2023; Wimber et al., 2015) to observe the extent to which preparation induces the reinstatement of overall perceptual information in each condition. Given the dissociations between attention and expectation observed during target processing (Gordon et al., 2019; Wyart et al., 2012) and the apparent commonalities reported during their top-down preparatory states (e.g. Battistoni et al., 2017), our overall hypothesis was that both manipulations would lead to the pre-activation of anticipated contents but through at least partially different neural mechanisms.

## Methods

Methods are reported in accordance with the Committee on Best Practices in Data Analysis and Sharing (COBIDAS) M/EEG (Pernet et al., 2019).

### 1. Data and code availability

Original code has been deposited at <u>Github</u> and is publicly available as of the date of submission. Results have been deposited at <u>OSF website</u>. Raw data are available online at <u>OpenNeuro</u>.

#### Participants

Forty-eight participants (mean age = 22.06, range = 18-31; 29 women, 18 men, 1 non-binary) from the University of Granada were recruited and received from 20 to 25 euros, depending on their performance. Two additional participants were discarded due to low behavioral accuracy (less than 80%) or excessive noise in the EEG (more than 20% discarded trials). They were all native Spanish speakers, right-handed with normal or corrected vision, and signed informed consent prior to participation. Besides, to comply with COVID-19 guidelines, the temperature of participants was measured upon arrival (always <37°C), they confirmed to have had no illness symptoms in the days prior to the experiment and wore a face mask during the whole session. Sample size was calculated to achieve a statistical power of 80% for an estimated small effect size (Cohen’s d = 0.3) and three independent variables (block x category x cueing). Using PANGEA (Power aNalysis for gEneral ANOVA designs(Westfall, 2016) we obtained a minimum of 32 participants to be able to detect the block x cueing interaction in reaction times and behavioral accuracy, our main behavioral prediction. To fit the counterbalancing scheme, we tested 48 participants. This sample size provides an estimated power of 94% under the described parameters.

### 2. Apparatus, stimuli, and procedure

Stimulus presentation and behavioral data collection were done with The Psychophysics Toolbox 3 (Brainard, 1997) on MATLAB (v.2020a) in a Microsoft PC. Stimuli were presented on an LCD screen (Benq, 1920×1080 resolution, 60 Hz refresh rate) over a grey background. We employed 160 male and female faces (50% each, with ~6°x9° visual angle, extracted from The Chicago Face Database(Ma et al., 2015)) plus 160 unique Spanish male and female names (50% each, with ~8°x2° visual angle). Four different geometrical shapes (circle, square, rain-drop and diamond with thin black outlines, unfilled, ~2°x2° visual angle) were used as cues in the main task. The sound stimuli employed in the localizer blocks consisted of four different tones (250, 300, 350 and 400 Hz).

The main task was a cue-target paradigm where, depending on the block, cues carried information about either the *relevance* (Attention) or the *probability* (Expectation) of upcoming face or word targets. Each trial started with the presentation of this visual cue. For each participant, and to avoid perceptual confounds, two cue shapes (counterbalanced across participants) were associated with faces and two with names. Importantly, cue pairs (the cue associated with faces and names) changed through the experiment. This way, the first cue for faces (e.g. a circle) appeared in half of the blocks with the first cue for names (e.g. a square) and the other half with the second cue for names (e.g. diamond). The task was to indicate the sex/gender of this target (male or female). Participants pressed one of two keys (“a”, “l”, counterbalanced across participants) to respond whether or not the target belonged to the gender stated at the beginning of each block. Half of the blocks belonged to the Attention condition, and the other half to the Expectation condition. Participants were verbally instructed to use the cues in the two blocks to respond as fast as possible while avoiding mistakes. At the beginning of each block (Figure 1B), they were informed about the block (Attention or Expectation), the target sex/gender (“Is the target male/female?”), and the two cues (one for faces and one for names). Importantly, and since Attention and Expectation are involved in almost any act of visual perception, we aimed at manipulating one process while keeping the other constant. In Attention blocks, the cue indicated the *relevant* stimulus category to select (either faces or names). Only if the stimulus belonged to the relevant category (50% trials, cued), the participant had to perform the gender discrimination task on the target. Otherwise, participants had to answer ‘*no*’ regardless of the stimulus sex/gender (non-relevant category, uncued). Note that this manipulation of relevance, where further processing has to be applied only to selected stimuli, is similar to that employed in previous literature (e.g. Baldauf & Desimone, 2014b; Saenz et al., 2002; Summerfield et al., 2006; Womelsdorf et al., 2006). Therefore, participants had to give an answer to all Attention trials, and had to be prepared to perform the gender judgment task. Importantly, both relevant and non-relevant targets were matched in expectation, as by design they appeared with a 50% probability after each attention cue. On the other hand, in Expectation blocks the cue indicated the *probable* category of the target, with a 75% likelihood (e.g.(de Lange et al., 2013; Kok et al., 2017) for similar manipulations). Here, participants had to perform the gender discrimination task in all trials, whether or not the target was cued. This way, both the expected and unexpected targets were equally relevant.

**Figure 1.**
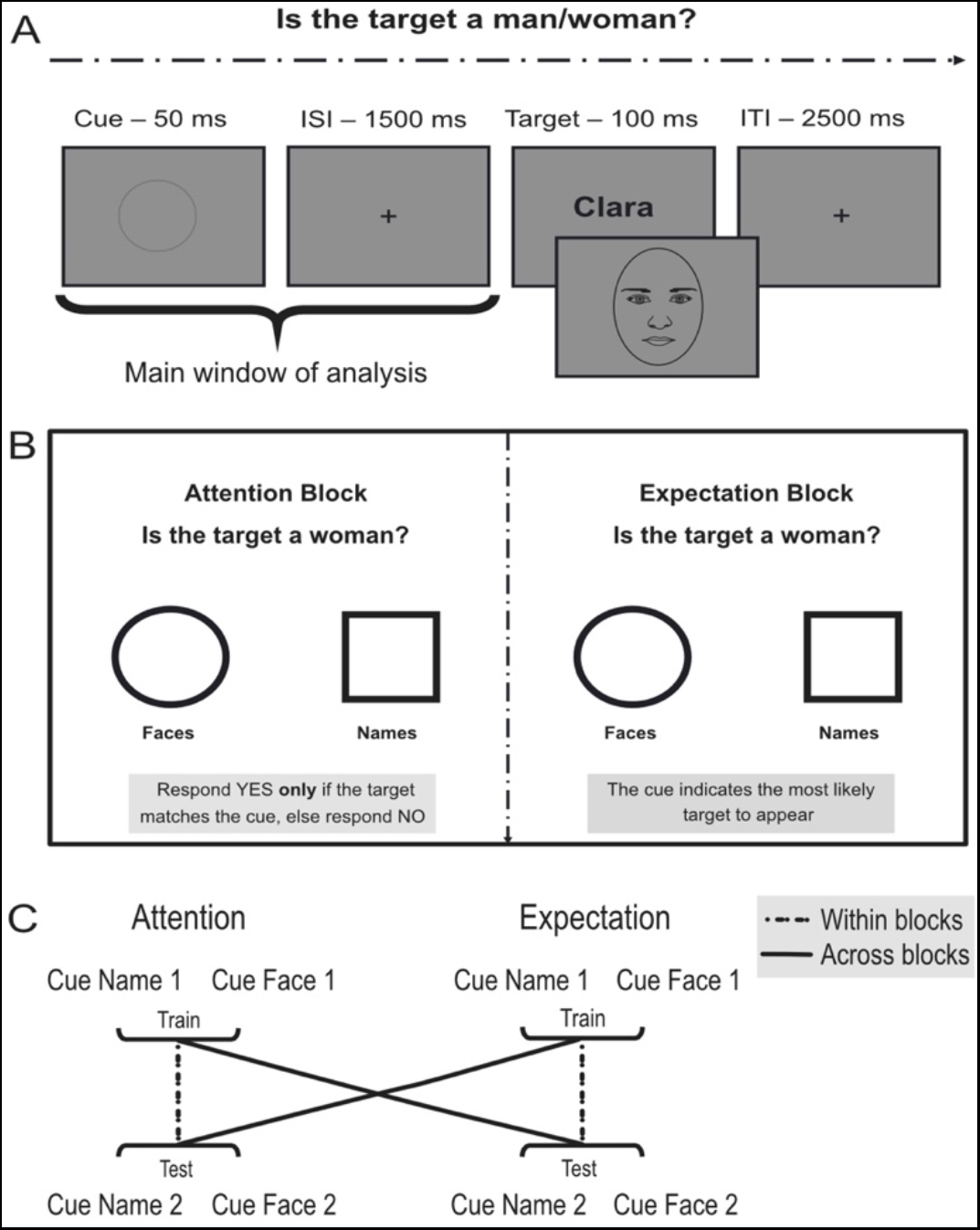
Behavioral task and design. **(A)** Behavioral task: example trial. Participants were cued about an incoming target stimulus (a face or a name) with which they performed a sex/gender classification task. **(B)** Block starting screen. In Attention blocks, the cue indicated the relevant stimulus. Participants performed the judgment on top of the screen only if the stimulus matched the cue. In non-relevant trials, participants responded with the “no” key. In Attention blocks the cues were not predictive of the probability of the target (50% faces vs. names) or the response. In Expectation blocks the cue indicated the probability of the stimulus category, which appeared 75% of times. Participants had to perform the judgment regardless of the cue (thus predicted and unpredicted targets were equally task-relevant), which carried no response information either. That is, in both blocks participants responded in all trials and perceptual details were fully equated. **(C)** MVPA classification rationale. In cue-locked within-block analyses, the classifier was trained to differentiate between anticipated (relevant or probable, depending on the block) faces vs. names with one pair of cues and then tested on the other, in either attention or expectation conditions. Note that classifiers could not use block information to differentiate between faces and words, as these were matched in every contrast. Across-blocks, the classifier was trained with one pair of cues within one condition (e.g., attention) and then tested on the other pair of cues within the other condition (e.g., expectation).

In every trial of the main task, the sequence of events was as follows: a 50 ms cue was followed by a fixed Cue-Target Interval (CTI) of 1500 ms and then the target appeared for 100 ms. Trials were separated by 2500 ms intervals. Auditory (tone, 400Hz, lasting for 300 ms) and simultaneous visual feedback (words “Attention” or “Expectation”, depending on the block presented for 500 ms) appeared in case of a wrong answer 1.3 seconds after target presentation, without altering the trials duration. Each trial lasted 4.15 seconds and each block 1.23 minutes. This main task was composed by 32 blocks of 20 trials each, or 640 trials in total.

In addition, Attention and Expectation blocks were interspersed with Localizer ones, used to measure target perceptual processing without overt motor activity (adapted from Egner et al., 2010). In these, the same stimuli as in the main task were presented. They were preceded by auditory cues that predicted either faces or names with 75% validity. Similar to the main task, there were two tones predicting faces, and two tones predicting names. One of each type was used per block. This manipulation was not used for the analyses reported in this work. Participants had to press a key *only* if the stimulus appeared upside down (rotated 180°, 10% of trials equated in stimulus category and sex/gender), regardless of cue validity. At the beginning of each block, a screen indicated the type of block and the two tones for the block. There were 16 localizer blocks of 40 trials each, 640 trials in total. The cue lasted 200 ms, the visual stimuli appeared after an ISI of 1500 ms and stayed onscreen for 100 ms. Trials were separated by 1500 ms intervals. Overall, each localizer trial lasted 3.3 seconds and each localizer block lasted 2.12 minutes.

The three types of blocks (Attention, Expectation and Localizer) appeared in a fully counterbalanced order, so that that they preceded and followed each other an equal number of times. Cues and target stimuli were also fully counterbalanced across participants. In total, the whole experimental session lasted approximately 80 minutes, with additional practice and EEG preparation time.

### 4. EEG data acquisition and preprocessing

#### 4.1. Acquisition

High-density EEG was recorded with 64 channels mounted on an elastic cap (actiCap Slim, BrainVision) at the Mind, Brain and Behavior Research Center (CIMCYC) of the University of Granada. Impedances were kept below 10 kΩ as recommended by the amplifiers’ manufacturers. EEG activity was referenced online to FCz and recorded at a sampling rate of 1000 Hz.

#### 4.2. Preprocessing

Data were preprocessed using EEGLAB (Delorme & Makeig, 2004) and custom MATLAB scripts (López-García et al., 2022). EEG recordings downsampled to 256Hz, digitally low-pass filtered using FIR filter at 120 Hz and high-pass filtered at 0.1 Hz. A notch bandpass was applied at 50 Hz and 100 Hz to remove line noise and its harmonics. Channels with excessive noise were identified by visual inspection and removed from the data (1.85% of channels on average, range 0-5). We used different epochs for cue and target stimuli and in each we split the data into 3 seconds epochs (−1 to 2 seconds after the onset of each stimulus). Then, Independent Component Analysis (ICA) was carried out to remove eye artifacts (i.e. blinks and lateral eye movements) using the *runica* algorithm from EEGLAB. Component rejection was guided by visual inspection of scalp maps, raw activity and power spectrum. ICLabel was used for further confirmation. In average 1.85 components were removed per participant (range 2-4). Next, we performed automatic trial rejection to prune the data from non-stereotypical artifacts. It was based on three factors: 1) abnormal spectra: trials in which the spectrum deviated from baseline by ±50 dB in the 0–2 Hz frequency window (sensitive to remaining eye artifacts) or deviated by −100 dB or +25 dB in 20–40Hz (sensitive to muscle activity); 2) improbable data: the probability of occurrence of each trial was computed by determining the probability distribution of values across trials, with a rejection threshold established at ±6 SD; (3) extreme values: all trials with amplitudes in any electrode out of a ±150μV range were automatically rejected (see (Keil et al., 2014; López-García et al., 2020, 2022) for similar preprocessing routines). The three methods in sum yielded an average of 8% of rejected trials per participant (range 1.8%-19%). Afterwards, the removed channels were repaired by spherical interpolation. We then applied common average to re-reference the data, given its widespread use and its optimal adaptation to high-density recordings(Pernet et al., 2018). Finally, trials were baseline corrected in the −200 to 0 ms prior to stimulus onset.

### 5. Analyses

#### 5.1. Behavioral

The main task design had three within-subject factors: Block type (Attention vs. Expectation), Cueing (Cued vs. Uncued) and Stimulus category (Faces vs. Names). We calculated three-way repeated measures ANOVA separately for behavioral accuracy and reaction times (RTs) employing JASP (Love et al., 2019). For each participant, trials with longer or shorter RTs than the average ± 2 SDs were discarded (11.54 % on average). Behavioral results in the localizer were not relevant and were just considered as exclusion criteria in case of poor performance.

#### 5.2. Representational Similarity Analysis

Representational Similarity Analysis (RSA) allows relating empirical multivariate measures of brain activity to theoretical models (Kriegeskorte et al., 2008). We performed RSA using voltage values on a subject-by-subject basis. Prior to the analyses data were normalized by z-scoring the values across all trials, regardless of the condition. We then constructed empirical Representational Dissimilarity Matrices (RDMs) every three time points, which measure the geometrical distances between all experimental conditions (see MVPA section) and, finally, estimated the relationship between empirical RDMs and theoretical models.

RDMs were built with data from eight conditions, yielding 8×8 symmetrical matrices. These conditions were all the possible combinations between the design variables: *cue prediction* (faces and names); *cue shape* (shapes 1 and 2 for faces, shapes 3 and 4 for names) and *block* (attention and expectation). We employed a Linear Discriminant Contrast (LDC, also known as Crossvalidated Mahalanobis Distance, (see Walther et al., 2016) as measure of distance between conditions, for the following reasons: it is a continuous measure, so it is highly reliable, informative, and lacks a ceiling effect; it includes a crossvalidation loop, which makes it less prone to biases; and it is centered around 0 when the true distance is 0 and therefore it is easier to interpret and more generalizable (Nili et al., 2014). We calculated LDC as described in Bueno & Cravo (2021). For every time point and each pair of conditions, we calculated the mean of each channel. This was done in two different datasets (train and test), to perform two-fold crossvalidation. We used two-fold crossvalidation to minimize computational costs, while avoiding biases due to random noise (Walther et al., 2016). The distance between the two conditions in the two folds was multiplied by the pseudo inverse covariance matrix between the residuals of the first and the second conditions in the training set, and the distance values were then averaged across the two folds.

Theoretical RDMs (Figure 3a) were built based on the expected distances (assigning values of 0 or 1) between conditions, according to different hypotheses. We built three such model RDMs based on: (1) cue shape (increased similarity between cues with same shape, regardless of the block and predicted target); (2) category (increased similarity between cues that predict the same stimulus category); and (3) block (increased similarity between cues belonging to the same block). The next step was to estimate the share of variance that each of these three model RDMs explained. To do so, at each time point we fitted a linear regression with the model RDMs as regressors and the empirical matrix as dependent variable. As a result, we obtained a t-value for each model, time point and participant that explained a significant *unique* portion of the variance, above that explained by the other regressors.

We used a non-parametric cluster-based permutation method to infer statistical significance at the group level, against empirical chance levels. First, at the single-subject level we randomly permuted the labels of the theoretical matrices. These permuted matrices were used as independent variables on a linear regression, which was repeated 100 times per participant. This gave 100 chance level t-values per participant and model. Then, one t-value for each model was randomly drawn per subject and the selected values were averaged. This was repeated 10^5^ times to obtain 10^5^ permuted group t-values. For each time point, the empirical chance distribution was estimated. As expected, this yielded a t-value distribution centered around 0. The above and below thresholds were estimated so that they included the 99% of the distribution. Groups of consecutive time points with values outside the previously calculated thresholds were measured generating the null distribution of cluster sizes. Finally, to further ensure correction for multiple comparisons, we used a False Discovery Rate (FDR) approach to determine the smallest cluster size deemed significant for α = 0.01 (López-García et al., 2022; Stelzer et al., 2013).

Besides, to estimate the topography of each model we repeated the analyses for each channel separately, estimating the empirical matrix using all time points. That is, at each channel we fitted a linear regression with the theoretical models as regressors.

#### 5.3. Time resolved Multivariate Pattern Analysis (MVPA)

We used time-resolved MVPA (Grootswagers et al., 2017) to study stimulus category-specific preparation by classifying faces vs. names. Note that we did not intend to directly classify attention vs. expectation, as this approach would be biased by existing block differences between these conditions. Instead, classifiers were trained and tested to differentiate between faces and names within blocks, and comparisons between attention and expectation were always performed with this base contrast. Also, classifications were done *before* stimulus presentation with cue-locked EEG, where faces and names were anticipated as relevant or probable by the cue (targets were not relevant for this work and thus not analyzed). The steps of the classification were equal for all analyses unless otherwise specified and were performed with voltage values. Classification was performed using MVPAlab (López-García et al., 2022) running on MATLAB. To maximize observations while reducing computational costs, we performed an MVPA classification every three time points. This way features used for classification were trials by channels matrices of raw voltage in single time points *t_n_*.

We applied two strategies to increase signal-to-noise ratio. First, the trials used were the result of averaging sets of three trials (randomly selected) in each condition (Grootswagers et al., 2017). Then, we employed a smoothing method, based on a moving average filter with a length of 3 time points. For every t_n_ the features of the previous and the following data points were averaged, so that t_n_ = (t_n-1_+ t_n_+t_n+1_)/3. Then, classes were balanced by subsampling the class with more trials so that the number of trials from the two conditions fed to the algorithm stayed the same (Grootswagers et al., 2017). A five-fold stratified cross-validation loop was implemented (King et al., 2013), which ensures that the proportion of each class stays balanced across folds, thus increasing the classification applicability to unknown data. Data were divided into five parts, which are enough to obtain unbiased results with a relatively small computational cost (Grootswagers et al., 2017; Varoquaux, 2018). The classification algorithm was trained with the first four divisions (training set) and tested on the remaining one (test set). This was repeated five times with the different sets. To improve the performance of the classifiers and the generalizability of multivariate analyses results (Singh & Singh, 2020), we normalized the data. Normalization was carried out within the cross-validation loop (King & Dehaene, 2014). Within each fold, we calculated the mean and standard deviation of each electrode across the training trials. The train set (X_train_) and testing set (X_test_) were normalized as follows:

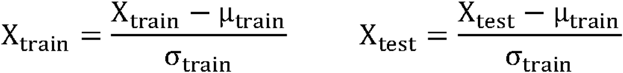

Where μ_train_ and σ_train_ are the mean and standard deviation of the training set.

We used Linear Discriminant Analysis as classification algorithm, given its good fit to typical EEG variability and higher sensitivity than similar methods (e.g. Support Vector Machines, see (Grootswagers et al., 2017; Kerrén et al., 2018a)). Time ranges were from −100 ms to 1550 ms. Classification results were estimated with an empirical receiver operative curve (ROC) analysis and reported as the area under the curve (AUC). This method works as an estimate of the true positive rate as a function of the false positive rate. The AUC is a non-parametric criterion-free method, so it does not involve assumptions about the true distribution of the data (King & Dehaene, 2014). Besides, it is less susceptible to systematic biases and it is especially sensitive to two-class differences. AUC results can be interpreted similarly to classification accuracy, with 0.5 indicating equal probability of true positives and false positives and 1 accounting for perfect discriminability between classes (King et al., 2013).

To estimate statistical significance, we again used cluster-based permutation analyses. In this case, the labels of each trial were randomly permuted. This was repeated 100 times per participant, generating chance level results. After following the same process as in the RSA section (using AUC instead of t-values) we ended up with a distribution centered around chance levels (0.5).

##### 5.3.1. Temporal Generalization Analyses

To characterize the changes of the signal throughout the temporal window, we employed a temporal generalization approach (King & Dehaene, 2017). On each time point we trained a classifier following the process described above. Then, we tested it on all time points of the preparation time window. This rendered a Temporal Generalization Matrix (TGM) representing the AUC values for each train-test pair. Statistical significance was then calculated for each TGM following the same rationale as with time-resolved analyses. The only difference was setting the minimum statistically significant threshold for cluster sizes to p<0.001, to avoid small clusters.

##### 5.3.2. Category-specific anticipation within attention and expectation contexts

The process described was applied to the cue-locked EEG separately for Attention and Expectation blocks, training classifiers to tell apart data from trials in which the cue anticipated faces vs. names. In a first approximation, we used trials of all cue shapes in each category (Figure 1c). Next, to ensure that classification results were not biased by perceptual differences between the geometrical shapes used as cues, we implemented a classification approach across cues. We first trained the classifier with data from only one pair of cues (e.g., classifying between squares predicting faces and diamonds anticipating names) and then tested it on the other pair (e.g., testing with circles that predicted faces and drops that predicted names). Because of the design, the selected training and testing pairs of cues only appeared together in half of the blocks. This cross-classification ensures that only the common differences between the two classified pairs will be decoded from the results (Kaplan et al., 2015), thus removing perceptual confounds. We averaged the results of both directions and their permutation maps to obtain a greater signal-to-noise ratio and to reduce biases due to the classification of specific perceptual features. The averaged results were then fed to the same statistical algorithm used previously to obtain cluster-based thresholds of statistical significance.

Once the results of these classifications were obtained, we compared the scores in Attention vs. Expectation blocks. To do so, we subtracted the empirical results of the two conditions (Attention – Expectation). Similarly, we subtracted the results of one of the 100 permutated chance level accuracy scores obtained during the cluster analysis in one condition from one from the other. We used the same cluster-based permutation implementation described above to evaluate differences in the two directions (Attention > or < Expectation). In this analysis the permuted distribution is centered around zero. Since we (arbitrarily) subtracted Attention - Expectation, positive values indicated greater results for Attention, while negative values indicated greater results for Expectation.

##### 5.3.3. Cross-classification between Attention and Expectation

To estimate the degree to which patterns of brain activity are shared for preparation across Attention and Expectation, we employed a cross-classification approach. We trained a classifier with data from one condition and then tested it on the other. We first did this using data from all trials in each condition, incorporating all four types of cues. Again, to rule out perceptual confounds we repeated the analyses following the same rationale described in the previous section (see Figure 1c for a summary of the condition selection strategies). In addition, we performed a control analysis to ascertain that cross-classification between Attention and Expectation blocks was feasible. Here we trained and tested using cues with the same physical form (e.g. train circle vs. square in attention, test circle vs. square in expectation) to observe whether the classifiers could extract the physical patterns of the cues even across overall changes in block demands.

#### 5.4. Canonical Template Tracking

Finally, we compared the sustained patterns that arose during the preparatory interval with the actual perception of face and name stimuli. To do so, we obtained Canonical Template Patterns (González-García et al., 2021b; Palenciano et al., 2023; Wimber et al., 2015) of brain activity generated by faces and names in the independent localizer blocks. First, we performed an MVPA analysis in localizer trials following the same process as in previous analysis. Then, we selected the time window where the classifiers locked to the localizer target stimuli in the localizer had the highest AUC (i.e., when activity patterns were more dissimilar) across participants (Figure S1), which was 100 – 300 ms after stimulus onset. Then, and separately for face and name localizer trials, we averaged the raw information of every time-point and trial in the selected window for each channel and category. This resulted in a vector of 64 channel activity values for faces and another one for names in each participant. Next, these CTP of faces and names were used as regressors in a linear regression where the dependent variable was the raw channel activity for each channel and condition during every time-point in the preparation or target window of the main experimental task. This rendered two t-values per time-point that accounted for the variance explained for each CTP (faces and names). We did this analysis separately in Attention and Expectation, for cues predicting faces and names and for face and name targets. To estimate statistical significance, we used the same cluster-based permutation analysis described above. Briefly, before averaging localizer data to create the templates for faces and names, we randomly permuted the trials of both conditions. This was repeated 100 times for each stimulus. Then, the randomly permuted CTPs were used as regressors, which gave 100 t-values for each participant and template. Then, the process was identical to the one we used in the RSA analysis. To estimate significance in when comparing the results for Attention and Expectation, we employed the method described in section 5.3.2.

Importantly, we used localizer data only from the targets, even though probabilistic cues were also included in these blocks. The localizer only included probabilistic (not relevance) cues, so it would not give equal insight into both relevance and probability contexts in the main task. Also, we considered that overall task demands between the localizer and the main experiment were too large to interpret unambiguously a potential lack of generalization of anticipation between these two contexts. Furthermore, the localizer was not designed to elicit reliable preparatory neural activity, which was hence not tested.

## Results

### 1. Behavioral

Forty-eight participants completed a cue-target paradigm where, depending on the block, cues carried information about either the *relevance* (Attention) or the *probability* (Expectation) of upcoming face or word targets (Figure 1A). Analysis of participants’ behavioral results showed that preparation for the incoming target stimuli affected performance. A three-way repeated measures ANOVA on reaction times (RTs) showed main effects of Block (F_47,1_=56.45, p<0.001, ηp2 = 0.54), Cueing (F_47,1_=5.59, p=0.022, ηp2 = 0.11) and Category (F_47,1_=50.52, p<0.001, ηp2 = 0.52). Overall, responses were faster in Attention (M = 569 ms, SD = 0.01) than Expectation (M = 598 ms, SD = 0.01) blocks. Cueing affected RTs differently depending on the block (Block*Cueing, F_47,1_=9.07, p=0.004, ηp2 = 0.16; Figure 2A). Post-hoc tests showed no effect of Cueing for Attention (t<1) whereas there was an effect for Expectation (t_47,1_=3.6, p<0.001, Cohen’s d = 0.52). Expected (cued) targets elicited faster responses on average (M = 592 ms, SD = 0.06) than unexpected (uncued) targets (M = 603 ms, SD = 0.07). Category also induced significant differences. Post-hoc tests showed faster responses for faces (M = 576 ms, SD = 0.7) than for names (M = 591 ms, SD = 0.75), t_47,1_=7.51, p<0.001, Cohen’s d = 1.08; Figure 2C.

**Figure 2.**
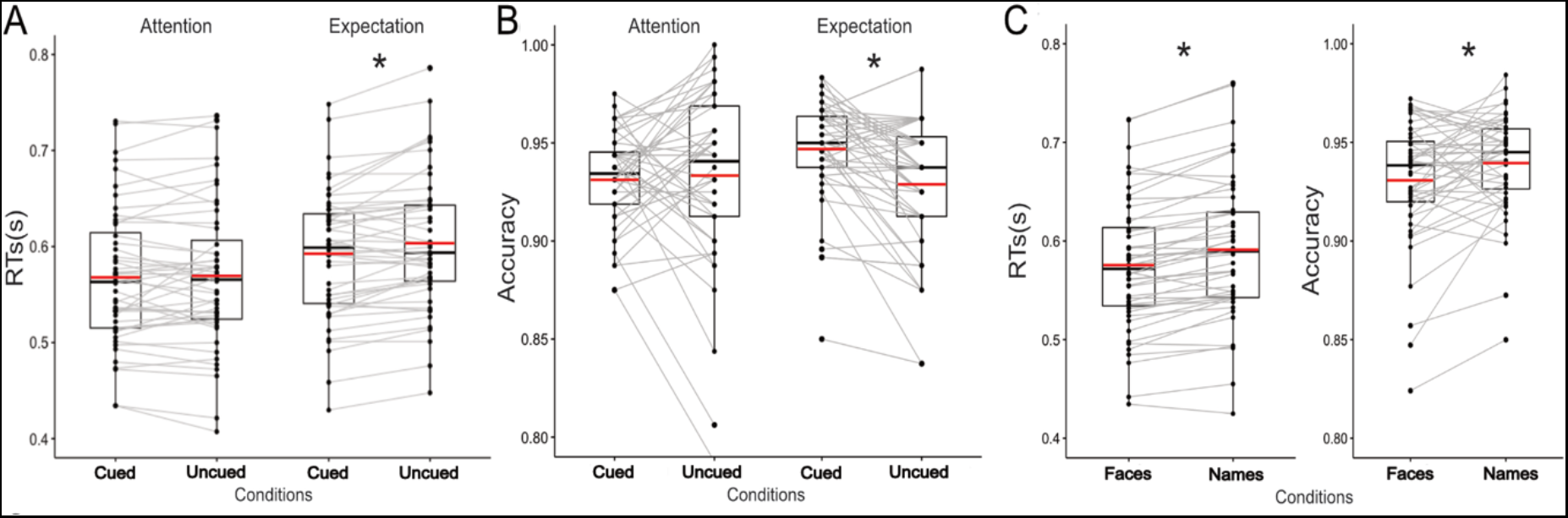
Behavioral results. Participants responded with a yes/no key press to the question asked at the beginning of each block. Note that in attention blocks, uncued trials were always responded with a “no” response. (A) Reaction times (in seconds) in Attention and Expectation blocks, for cued and uncued trials. (B) Accuracy in Attention and Expectation blocks, for cued and uncued trials. (C) RTs and accuracy for face and name targets, averaged across conditions. Dots represent individual subject (n=48) scores per experimental condition. Grey lines connect each participant’s score in the two conditions of each block. The horizontal black line inside boxes represents the median, the horizontal red line represents the mean, while the limits of the box indicate the first and third quartile. Note that due to the high SD, the mean and the median may differ slightly. Whiskers indicate the 1.5 inter quartile range for the upper and lower quartiles.

The ANOVA on behavioral accuracy showed results in the same direction albeit less prominent. Both Cueing (F_47,1_=4.33, p=0.043, ηp2 = 0.01) and Category (F_47,1_=7.04, p=0.011, ηp2 = 0.011) were significant, and there was no main effect of Block (p>0.05), with both conditions presenting high accuracy overall (Attention: M = 0.92, SD = 0.07; Expectation: M = 0.93, SD = 0.07). Again, we found that cueing affected each block differently (Block*Validity, F_47,1_=9.67, p=0.003, ηp2 = 0.15; Figure 2B). Although there were no differences in Attention (t<1), there was an effect of expectations (t_47,1_=3.59, p=0.003, Cohen’s d = 0.52), with cued trials eliciting more accurate responses (M = 0.947, SD = 0.026) than uncued ones (0.929, SD = 0.035). Post-hoc tests showed less accurate responses to faces (M = 0.93, SD = 0.4) than to names (M = 0.94, SD = 0.37; Figure 2C), t_47,1_=2.65, p=0.011, Cohen’s d = 0.38; Figure 2C. Overall, these results indicate that, as instructed, cues were used effectively and also differently across blocks.

### 2. Time-resolved profile of preparation

Our first aim was to assess the emergence of specific coding patterns linked to different information content during preparation in Attention vs. Expectation contexts. The perceptual features of the cue should be processed, and their contextual meaning extracted to anticipate general target-category information. This anticipated target category should be activated and maintained in working memory to later provide an efficient response. To assess the contribution of each of these sources of information to preparatory activity we employed model-based RSA (Kriegeskorte, 2008).

Results of the multiple regression for the three RSA models built (Cue shape, Category and Block, see Methods section and Figure 3A) revealed the unique variance explained by each of the factors entered in the analysis. A peak of the Cue shape model appeared first, at 160 ms after cue presentation, and decayed fast afterwards (Figure 3B, red line). In contrast, the coding of the specific incoming target category increased progressively along the interval, reaching its peak right before the presentation of the actual target (Figure 3B, green line). In addition, the variance explained by the blocks reached its peak around 500 ms after cue onset, slowly decayed along the interval, and fell below significance levels 100 ms before target onset (Figure 3B., blue line).

**Figure 3.**
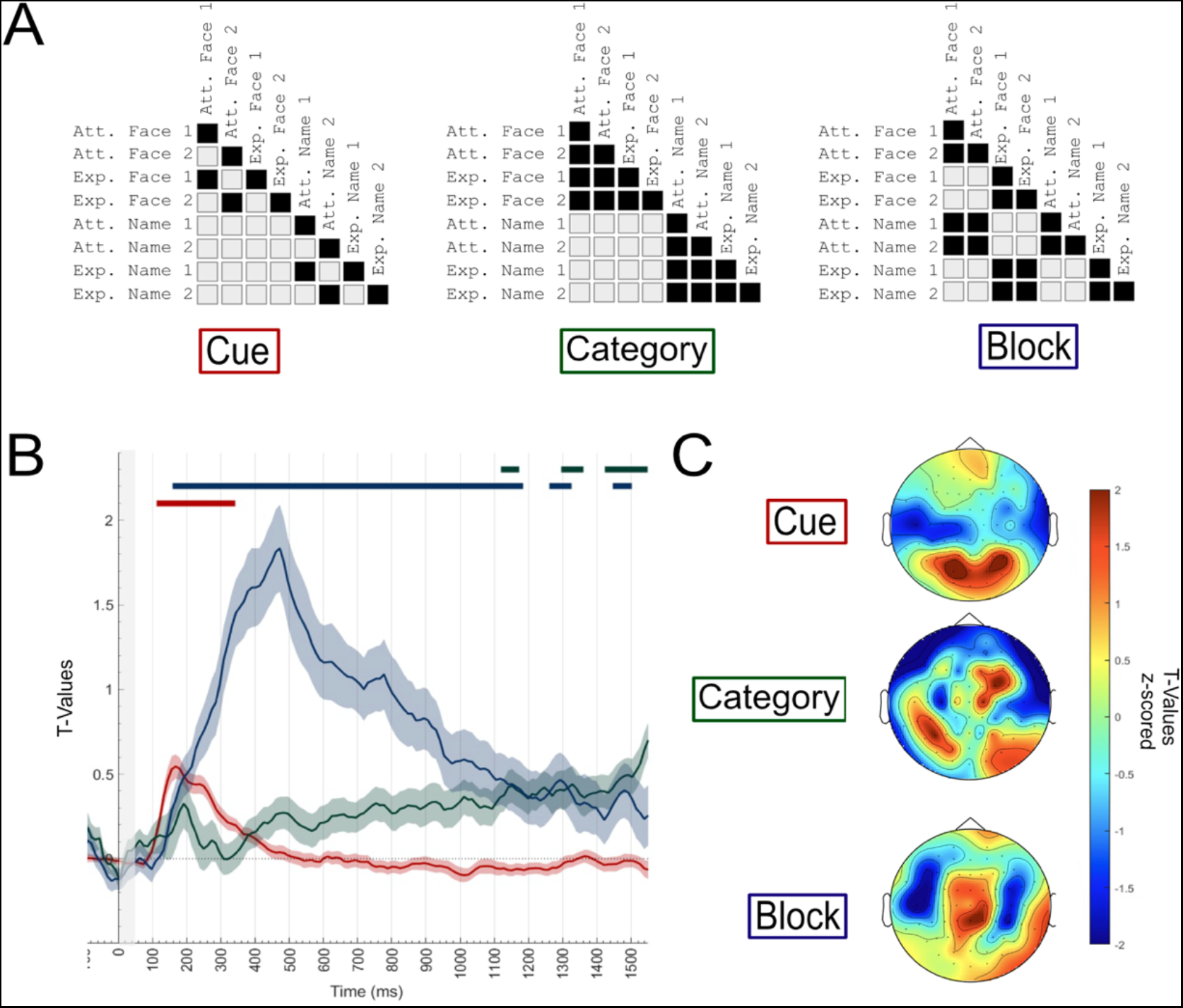
(A) RSA theoretical matrices and (B) results. In (A), black squares represent high similarity between conditions and grey squares represent low similarity. In (B), each line depicts t-values from a linear multiple regression fit for the three theoretical dissimilarity models, showing the unique share of variance explained for each of the factors. The colored straight lines above indicate significance clusters after a cluster-based permutation analysis. Grey shading indicates cue presence onscreen. The cue model (red) indicates dissimilarity between cue shapes, regardless of the block or prediction; the category model (green) indicates similarity between cues predicting faces or names; the Block model (blue) shows overall preparatory differences between Attention and Expectation. (C) Topographies of each model after using time points as features and repeating the analysis on each EEG channel.

### 3. Time-resolved classification shows increasing AUC throughout the preparation interval

We then studied the coding of the anticipated specific information during the preparation interval, separately for Attention and Expectation. Importantly, we did not compare cued vs. uncued targets, but cues that predicted (relevant or probable, depending on the block) face vs. word stimuli within the same block. Using the two cues of each category together for training and testing returned a classification weighted on the perceptual features of the cues (see Figure S2 for a detailed description of the result), similar to the Cue model in Figure 3 (red line). Afterwards, we employed a cross-classification analysis between different cues to avoid cue perceptual confounds in the classification (Figure 4). Since two differently shaped cues coded for each type of category, we trained the classifier in one pair and tested it on the other, repeated the process in the opposite direction and averaged both. Similar to the Category model in Figure 3, time resolved cross-classification showed that the accuracy in the decoding of the anticipated category increased as the target onset approached. We employed the subtraction approach described in the methods section, but we found no differences in classification between Attention and Expectation (all ps>0.05). We repeated a regression analysis to predict the cross-classification result of each participant based on the average time-resolved result (t-values) of each RSA model. As expected, the category model (Figure 3, M = 0.35, SD = 2.37) explained the results better than any other. T-values were nominally higher than the cue model (M = 1.04, SD = 1.42) and significantly higher than the block model (p<0.01).

**Figure 4.**
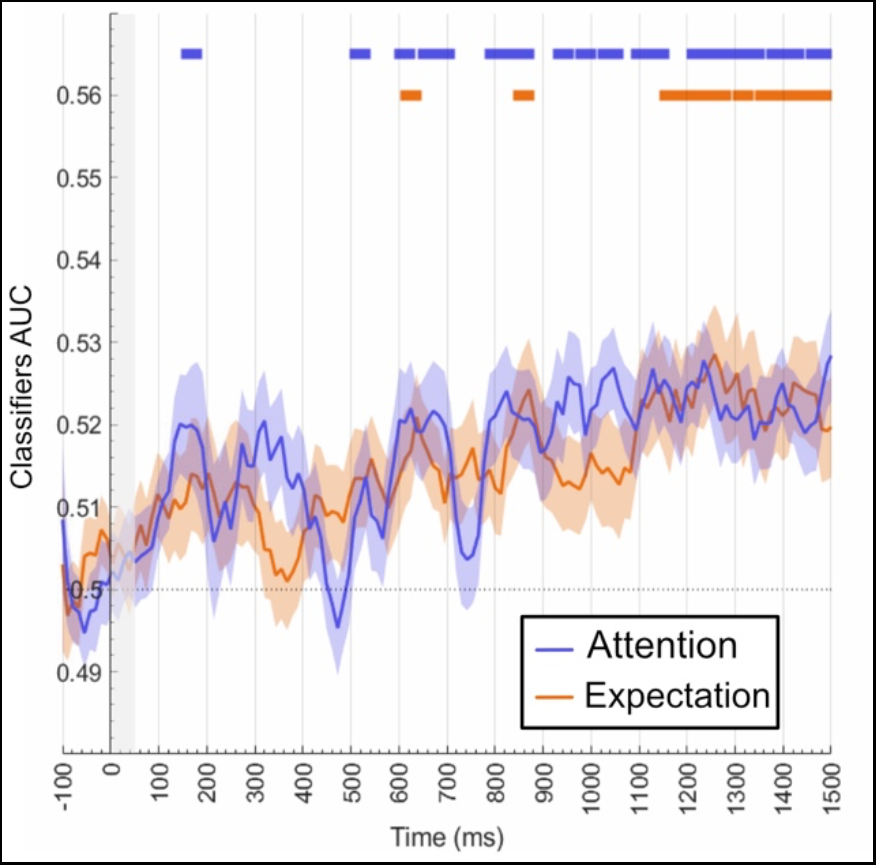
Raw voltage time-resolved cue decoding results. Result of the time-resolved classification of the category (faces vs. names) to be selected (blue) vs. expected (orange). Horizontal colored lines indicate statistical significance against chance within each block. Grey shading indicates cue presence onscreen. Figure shows the result of training and testing as described in Figure 1C (dotted line). For the results using all the cues together for training and testing see Figure S2.

### 4. Relevance- and probability-driven preactivations are stable

While the previous results provide an initial characterization of neural coding of specific category information during the preparation interval, they do not allow to explore the extent to which relevant representations are stable during this time window, since significant decoding on different time points could be driven in principle by different mechanisms. To investigate this, we employed a cross-time decoding approach (King & Dehaene, 2014) to compare different patterns of brain activity across the preparation interval. For this, we trained a classifier in one time point and then tested it on all the points of the interval (Figure 5A, B). Results showed clear signs of generalization during the preparation interval in both Attention and Expectation conditions (black outlines). The clusters of activity grew increasingly larger up until the target’s onset, indicating that the underlying patterns remain relatively stable during the preparation period. We then compared the results of both analyses following the same rationale as above: we subtracted the result matrices for both conditions and performed a one-side T-test against 0. This analysis did not yield any significant results, indicating that the accuracy of anticipatory category decoding was not different between conditions. Altogether, these results suggest that preparation in both Attention and Expectation leads to a similar level of discriminability of the anticipated (relevant or probable) category. However, this analysis is agnostic regarding potential similarities in how anticipated relevant vs. probable information is coded, as different underlying mechanisms could lead to similar accuracy results.

**Figure 5.**
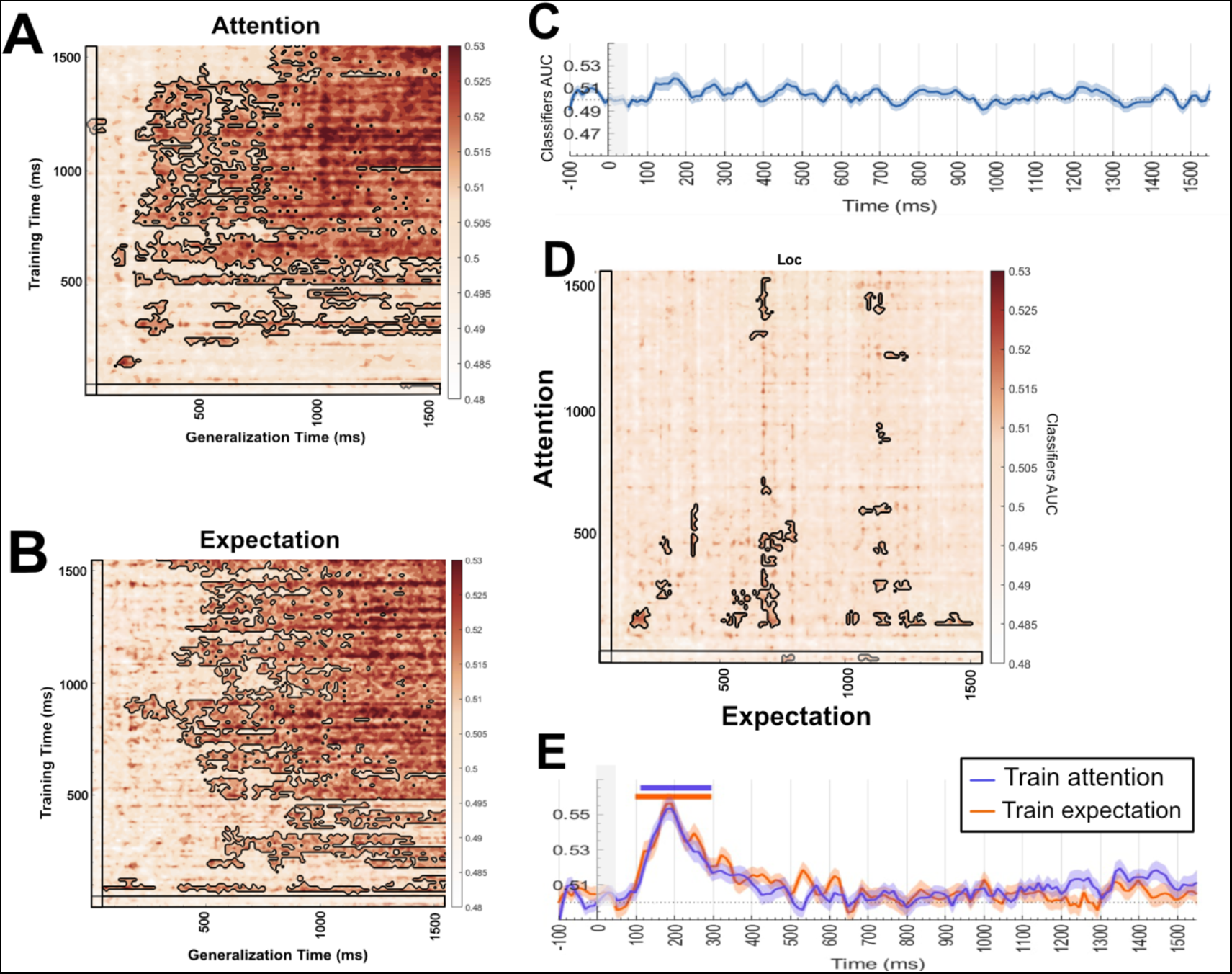
Raw voltage temporal generalization results. (**A-B**) Temporal generalization results using the cross-cue decoding scheme, within (**A**) Attention and (**B**) Expectation conditions. Black outlines indicate statistical significance against chance. Black boxes indicate cue presence on screen. **(C-D)** Cross-condition classification results. Classification using the scheme described in Fig 1C, depicting the average of the two train and test directions. **(C)** Time-resolved classification of anticipatory information coding after training and testing the classifier in different conditions yielded no significant decoding. **(D)** Temporal generalization matrix. For visualization purposes, we averaged training and testing in the two directions (train attention and train expectation). The horizontal axis shows the times for training and testing in Expectation, and the vertical axis that of training and testing in Attention. That is, we inverted the axis of one of the directions before averaging. We chose this rationale because activity that is better represented in only one of the conditions should appear only on one side of the diagonal (e.g., above) when training and the opposite when testing (e.g., below). Averaging this way, we can show information that would not be visible doing a standard average. The non-averaged results for both directions can be found in the Supplementary Section (Figure S1). **(E)** A control cross-classification between Attention and Expectation (*Orange: Train Attention – Test Expectation. Purple: Train Expectation – Test Attention*) employing the same pairs of cues to train and test the classifier (e.g. train with circles in Attention, test with circles in Expectation) shows that classification across blocks is feasible. However, whereas the perceptual features of the cues generalize across Attention and Expectation (significance is marked with colored bars), the specific preactivation of contents before target onset does not.

### 5. Attention and Expectation induce distinct patterns of preparatory activity

To examine the mechanisms supporting the classification results, we used multivariate cross-classification by training the classifier in one condition and then testing it on the other. Similar patterns of brain activity should generalize between conditions, while differences should provide chance-level classification. The number of observations employed for this, as well as the inter-block temporal distance, were also matched. The averaged results (training in attention and training in expectation) for raw voltage cross-classification are shown in Figures 5C, D (see Figure S3 for the results split by train and test direction). Common coding between anticipating relevance vs. probability of stimulus categories was scarce. Small significant clusters appear scattered through the temporal generalization interval. This result complements the previous analysis by suggesting that different neural mechanisms support the classification results. In addition, the control analyses performed using the same physical cues to cross-classify between attention and expectation (see Figure 5E) show that generalization across blocks is feasible, given that the classifier shows significant above-chance performance to predict the perceptual shape of cues across attention and expectation conditions.

### 6. Tracking perceptual patterns of brain activity

We have shown that although Attention and Expectation lead to similar degrees of anticipatory classification, their underlying neural patterns are partially different. We hypothesized that such differences could arise due to the extent to which anticipatory representations function as perceptual reinstatements of the prepared target categories. In this case, the preparatory patterns that allow to classify anticipated faces and names should be similar to those triggered during actual stimulus perception. To assess such reinstatement of target representations during preparation, we obtained canonical template patterns (CTP; Palenciano et al., *2023)* of brain activity associated with faces and names (as described in the Methods section). We then fit a linear regression with the two CTPs as regressors to measure the extent to which these explained variance in preparatory activity across conditions, in each time point (see Figure 6).

**Figure 6.**
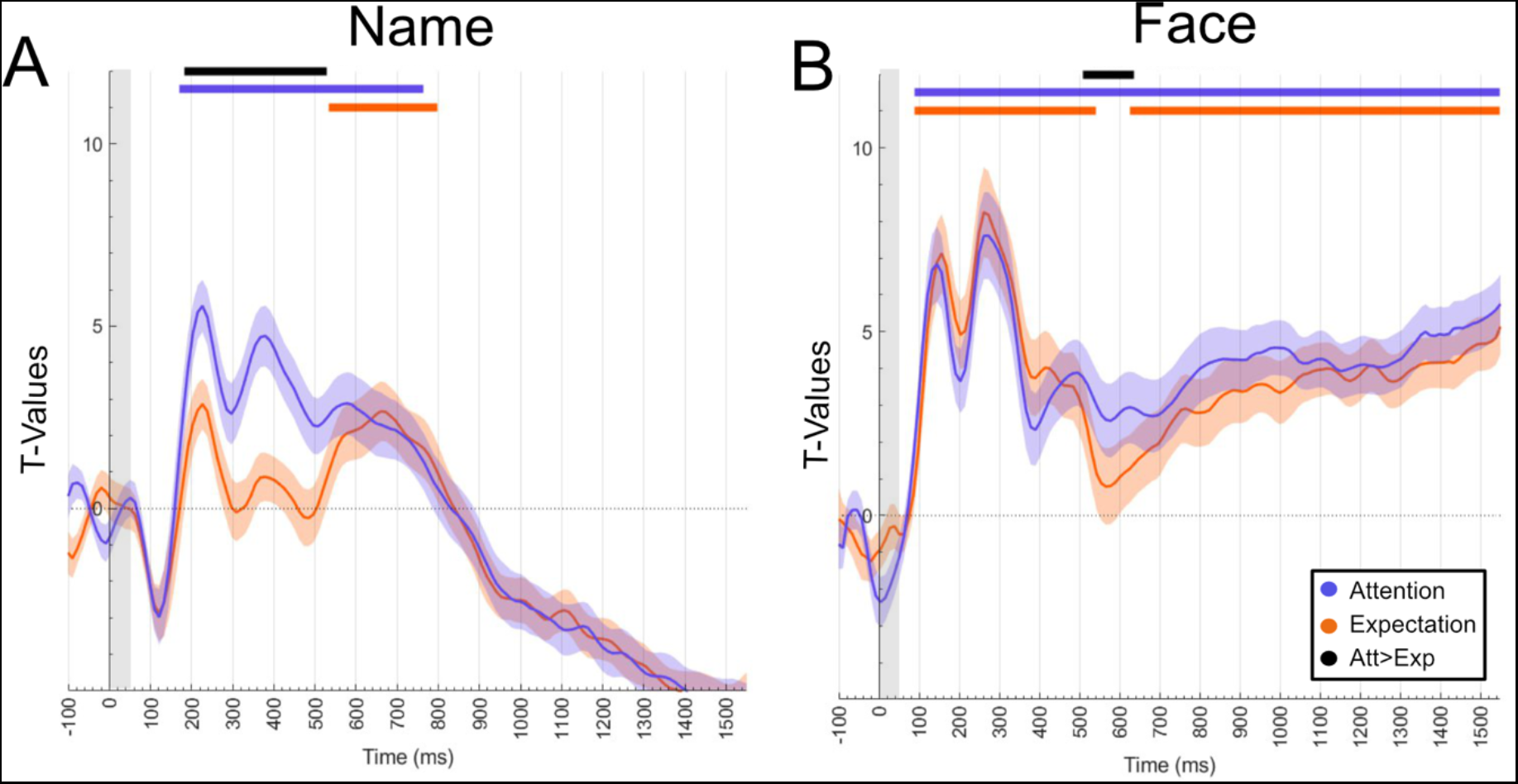
Canonical Template Tracking results. Grey shading represents cue on screen. **(A-B)** CTP results for separate cues for Attention and Expectation. Average of face and name cues **(A)** Name CTP model for cues. **(B)** Face CTP for cues. Horizontal lines depict significance. Black horizontal lines indicate points where the results for Attention were significantly higher than for Expectation. There were no clusters were Expectation showed higher reinstatement.

First, to complement the behavioral results regarding face and name processing, we applied the CTP to EEG activity locked to target onset in Attention and Expectation (Figure S4). The face CTP explained better the perception of target faces, while same was true for the name pattern on name perception. Moreover, face processing was more pronounced, but decreased more rapidly. Name processing, on the other hand, reached a lower peak but it decreased more slowly over time. As expected, these results further suggest that faces and names are processed differently.

Then, we compared how these canonical templates overlapped with preparatory activity in Attention vs. Expectation contexts. Since both CTPs similarly explained face and name predicting cues, here we show the average result of the two types of cues separately in Attention and Expectation blocks. Note that whereas such averaging prevents drawing any conclusion regarding selective preparation, it allows contrasting the extent of overall perceptual reinstatement across relevance vs. probability anticipation. Crucially, both perceptual templates were better predictors of category anticipation in the Attention compared to the Expectation condition (see Figure 6). Overall, these results indicate that regardless of the intrinsic differences in face and name processing, there is an overall stronger perceptual reinstatement for Attention than for Expectation. Note, importantly, that this stronger coding cannot be explained by overall increased difficulty, as responses were faster in attention compared to expectation blocks (see Behavioral Results section).

## Discussion

In the present work we studied and contrasted, for the first time, the top-down neural mechanisms engaged during category-specific preparation in two major information-processing contexts: Selective Attention and Perceptual Expectations. Our results reveal that the anticipated stimulus category is active during preparation, regardless of the specific demands. Crucially, we demonstrate that relevance and probability influence the preparatory patterns of brain activity in a unique manner, potentially via differences in the perceptual reinstatement of anticipated information. Overall, these results have important implications for theoretical models that explain how the brain anticipates relevant vs. probable forthcoming information.

Our behavioral results show that the paradigm manipulated Attention (relevance) and Expectation (probability) in an effective manner and as expected (Baldauf & Desimone, 2014b; Kok et al., 2017). Cued trials in Attention required further processing than those uncued, as participants were required to evaluate the sex/gender of the cued stimulus category and skip this judgment in uncued ones (Baldauf & Desimone, 2014a; Saenz et al., 2002; Summerfield et al., 2006; Womelsdorf et al., 2006). Importantly, for Attention the probability of either category was equal (50%), keeping expectation constant for relevant and irrelevant trials. The high accuracy in Attention blocks indicates that the cues were used effectively to select when needed and to respond. On the other hand, both expected and unexpected trials required selecting the forthcoming target, and thus equated relevance while leading to predicted and unpredicted stimulus categories. In line with validity effects repeatedly reported in the literature in expectation contexts, the efficiency of behavior increased in cued trials (de Lange et al., 2018; Sy et al., 2014). It may be important to note that our results are at odds with the notion that expected stimuli gain higher task-relevance and thus receive more attention that unexpected ones (see Rungratsameetaweemana & Serences, 2019), as this interpretation would lead to expect similar underlying anticipatory patterns for our attention and expectation manipulations.

We employed RSA (Carlson et al., 2019; Kriegeskorte, 2008) combined with multiple regression to study the temporal profile of information coding throughout the preparatory interval. This showed that anticipatory cues triggered several time-overlapping, yet distinguishable, effects in time (Figure 3B) and spatial arrangement (Figure 3C). The perceptual characteristics of the cue were coded early in the time interval. Immediately after, and during the majority of the interval and until around 100 ms before target onset, mechanisms pertaining to contexts capitalizing on either selective attention or expectations were deployed, as revealed by the Block model temporal profile. Evidence of anticipatory coding of different task sets has been found before (González-García et al., 2021a; Hebart et al., 2018; Ana F Palenciano et al., 2018). Interestingly, this result cannot be explained by motor preparation, since cues did not predict motor responses; instead, it is most likely due to the control context in each scenario. Finally, the variance explained uniquely by the Category model on anticipatory activity increased steadily and peaked before target onset. This pattern of emerging, ramping-up category representations could reflect two non-exclusive anticipatory mechanisms. Once the meaning of the cue is extracted and stored in working memory, the actual anticipation of a stimulus could be facilitated by top-down preactivation of perceptual regions (Auksztulewicz & Friston, 2016b; Trapp et al., 2016). Simultaneously, the effect of temporal expectations (as the preparatory interval was fixed) could induce an increasing preactivation of perceptual regions as the stimulus onset approaches (Jin et al., 2020; Rohenkohl et al., 2012). In either case, these results suggest that preparation engages a series of mechanisms that act both sequentially and in parallel, including the processing of bottom-up signals that are transformed into top-down category anticipation in a context of relevance or probability.

The block model in the RSA suggested that Attention and Expectation led to distinguishable coding patterns. One mechanistic explanation for this result is that the patterns for the anticipated category are more robustly coded in one condition over the other. Hence, we employed an MVPA approach (Grootswagers et al., 2017; Haxby et al., 2014) to classify the prepared category. Crucially, classifiers did not directly compare Attention vs. Expectation blocks (a discrimination that would be biased by differential block characteristics) nor cued vs. uncued targets (which would mix reactions to matched vs. mismatched predictions). Instead, classifiers were trained and tested to differentiate between cues that predicted (relevant or probable) faces vs. names, within each block. Results matched the RSA model, pointing to a robust effect of category anticipation. Furthermore, this prediction increased towards the end of the interval. Again, this resonates with literature on temporal anticipation (Barbosa et al., 2020; Jin et al., 2020; Ruz & Nobre, 2008). Moreover, prominent theories of attention (Mongillo et al., 2008; Trübutschek et al., 2017) pose that information can be held in WM by strengthening the synaptic weights between neurons, allowing for shifts (in this case, steady increases) in decoding results during the delay period without losing information about the maintained stimulus. The results revealed no significant differences in AUC between the conditions in either time-resolved or temporal generalization matrices, and suggest that anticipated targets are coded with similar fidelity during Expectation and Attention. Importantly, these results were obtained cross-classifying different pairs of cues, thus avoiding perceptual confounds. However, a similar degree of accuracy classification does not warrant identical underlying mechanisms. To test this, we performed multivariate cross-classification between Attention and Expectation. Surprisingly, despite well above-chance decoding within each type of block, there was little cross-classification between conditions. Importantly, classification scores were similar when training and testing in different blocks of the same condition (see Figure S5). In addition, a control analysis using the same cues showed that generalization across blocks was possible and not prevented by the differential block context. Classifiers trained and tested in different blocks were able to discriminate the perceptual nature of the cues, whereas the anticipated category cross-classification was absent (see Fig. 5). Overall, this set of results suggests that whereas Attention and Expectation both lead to anticipatory category representations, their top-down mechanisms are partially different, providing further support to the dissociation of relevance and probability, reported by previous studies, during target processing (Summerfield & Egner, 2009; Wyart et al., 2012a).

One possible explanation for the differences found is the degree of preparatory perceptual reinstatement (Kerrén et al., 2018b; Kilner et al., 2007; Muckli et al., 2015; Rose et al., 2016; Smith & Muckli, 2010; Vetter et al., 2014) in Attention and Expectation. We employed a Canonical Template Tracking procedure (González-García et al., 2021a; Wimber et al., 2015,Palenciano et al., 2023) to compare perceptual reinstatement in attention and expectation. We obtained canonical representations of the two target categories (faces and names) from an independent localizer, and then estimated the variance explained by these perceptual patterns during the anticipatory window. First, we applied the extracted canonical templates to the actual target processing, with results supporting that faces are processed differently from words. Importantly, when applied to anticipatory activity, we found that the canonical templates explained preparatory variance equally well for both predicted categories, but the reinstatement was significantly higher in Attention than in Expectation. Note that this cannot be due to a higher difficulty of the attention blocks, as RT showed the opposite pattern (faster responses in Attention). Instead, this higher reinstatement may be due to Attention directing more resources to activate perceptual codes in anticipation. It is unclear, however, why this analysis did not provide evidence of category-specific reinstatement, although it could be related to the mixture of perceptual activity caused by the cues themselves. It is possible that the physical shape of faces and words has differential overlap with the overall shape of the cues employed in the main task, which could have added additional variance to the overall analysis. This could have mixed with the anticipated category information and generated the lack of specificity. Relatedly, the patterns captured by our CTP may be different from those involved in categorical specific anticipation, which may have happened at a different abstraction level. Note that the large overall task differences between the localizer and the main task prevent the use of localizer cues to predict preparation during the main task. Further research employing different approaches to measure reinstatement should be conducted to clarify this matter.

How Attention and Expectation affect perception is an ongoing debate. Several studies have used paradigms that combine both processes to study how they interact. Attention has been suggested to sharpen the differences between expected and unexpected stimuli (Jiang et al., 2013), possibly changing the oscillatory profile of relevant categories (Auksztulewicz et al., 2017) while reversing repetition suppression (Kok et al., 2012). Although it has been suggested that attention acts from early processing stages, results so far are not conclusive (Alilović et al., 2019). Relatedly, Attention boosts signal-to-noise processing by suppressing noise, while Expectation increases baseline activity in perceptual regions (Wyart et al., 2012; see also Gordon et al., 2019; Rungratsameetaweemana & Serences, 2019). Predictive coding accounts propose that Attention increases prediction error of selected stimuli by suppressing noise of unattended categories, while expectation increases global sensitivity through prediction signals. Crucially, our results extend this literature by showing anticipatory differences between conditions, which cannot be accounted for by prediction error differences as targets have not been processed yet. Speculatively, attention could bias anticipatory neuronal sensitivity by increasing perceptual differences between relevant categories, coding these changes at least partially in the gamma band perhaps through anticipatory biasing of error processing units. Expectation could increase sensitivity to probable categories by increasing excitability of, perhaps, perceptual units.

Although overall our results are a crucial first step to characterize mechanisms across relevance and probability anticipation, they should be complemented by further studies. Although the results are statistically significant after robust cluster-based correction, accuracy values are lower than those obtained, for example, using target-locked data (e.g., Figure S1). Importantly, decoding accuracies do not equal effect sizes (Hebart & Baker, 2018). The values we obtained are within typical classification ranges when studying subtle neural patterns (Christophel et al., 2015; Hebart & Baker, 2018; also see Rose et al., 2016) employing non-invasive human neuroimaging. Future studies may obtain higher accuracies by increasing stimulus repetition and thus reinforcing specific neural traces of the information anticipated.

Moreover, it could be argued that the diverging results between the two conditions shown here are not due to differences in stimulus representation for relevance vs. probability, but to different task demands of the two blocks. However, as the goal of the study was not to fully differentiate the neural mechanisms of attention and expectation but to understand how anticipatory information is coded differently in contexts of relevance vs. probability, we did not merely contrast the two blocks directly. Instead, the analyses contrasted anticipated faces vs. words in conditions matched within blocks. In the same line, the arguably potential higher relevance of probable contents could not drive results, as such increased relevance would be equal for predicted probable faces vs. names, and thus the classifier cannot rely on this information. Moreover, and to further show how processes that are similar do generalize across both blocks, we trained the classifier in a pair of cues in one block, and tested it in the same pair for the opposite block (see Figure 5E). As expected, significant classification appeared for the cue identity, showing that cross-classification across blocks is feasible. If differential task demands had an overall effect changing the format of coding during anticipation, this should have arguably also altered cue patterns, preventing cross-classification. The fact that classifiers were able to extrapolate, though, adds to the idea that block differences do not account for lack of cross-classification of anticipated relevant vs. probable information content. Admittedly, classifiers can be conceived as black boxes where disentangling which factors are driving the results can be hard. However, the results obtained when employing RSA provide concurring evidence. Here, theoretical matrices (Cue, Category and Block) were used to study how these factors explain unique variance during the preparation window. Results (see Figure 3) suggest that Block (task demands) explains most of the variance during an earlier time window than Category, which in turn incrementally explains variance (and with a different topography from that of Block) as target onset approaches. Of note, the timing of this effect is quite similar to the window where classifiers show specific preparation for relevance and probability that does not cross-classify across these contexts. In any case, studies employing improved paradigms should be tested to replicate and validate these results.

Our scope was limited to the temporal domain, and questions arise regarding potential differences between brain regions. Internal predictions have been generally associated with the hippocampus (Aitken & Kok, 2022; de Lange et al., 2018; Hindy et al., 2016; Stachenfeld et al., 2017), which has a location that challenges EEG sensitivity. Further studies should employ more spatially sensitive techniques. Additionally, we focused on category-based preparation of faces and words, to facilitate having the same task across categories (sex/gender judgments). These two types of stimuli, although frequently combined in the literature (Alm et al., 2016; Amado et al., 2018; Dumas, 2015; Rose et al., 2016; Sperling et al., 2003) have a different spatial layout, which may have generated the anticipation of different spatial templates. Although this is not a confound in our task, as such difference is constant, it may have added a spatial component to the preparation. Finally, we focused our design on visual perception. Studies that have compared the effects of attention and expectation have used auditory and visual stimuli showing promising results (e.g. Jiang et al., 2013). It is likely that the mechanisms found here can be extended to other sensory modalities, but more research is needed to assess this idea.

Altogether, our results show that anticipatory representations are category-specific and also task dependent, and highlight the role of proactive processes that precede stimulus perception. These findings have important implications for current models of brain functioning. We show that instead of single, unitary top-down phenomena, the brain implements distinct modes of content-specific anticipation, which are tailored to task context and involve different levels of overall perceptual reinstatement. Predictive coding and attention models that differentiate these processes during target processing should be extended to the preparatory interval, acknowledging the specificity and complexity of these top-down anticipatory phenomena.

## Acknowledgments

This research was supported by the Spanish Ministry of Universities under the PID2019-111187GB-100 grant to MR and by the Spanish Ministry of Economy and Business under the TEC2015-64718-R to JG. JMGP is supported by a scholarship from the Spanish Ministry of Universities (FPU18/01853). CGG was funded by the Spanish Ministry of Science and Innovation (Grants Ref.: IJC2019-040208-I and PID2020-116342GA-I00), and Grant RYC2021-033536-I funded by MCIN/AEI/10.13039/501100011033 and by the European Union NextGenerationEU/PRTR.

## Author Contributions

Conceptualization, JMGP and MR; Data Collection, JMGP, DLG, and BAL; Analyses, JMGP and DLG; Resources, JG and MR; Writing – Original Draft, JMGP; Writing – Review & Editing, CGG and MR; Supervision, CGG and MR; Project Administration, MR; Funding Acquisition, MR

## Declaration of Interests

The authors declare no competing interests.

## Notes

https://osf.io/2rhjn/?view_only=1a729284b9d549b082ef725bc5081f3d

https://github.com/ChemaGP-UGR/AttExpLoc_EEG

https://openneuro.org/datasets/ds004502

### Competing Interest Statement

The authors have declared no competing interest.

### Summary of Updates

Added link to openly available raw EEG data.

