## Supplementary Materials for "Top-down specific preparatory activations for Selective Attention and Perceptual Expectations"

### Supplementary Section

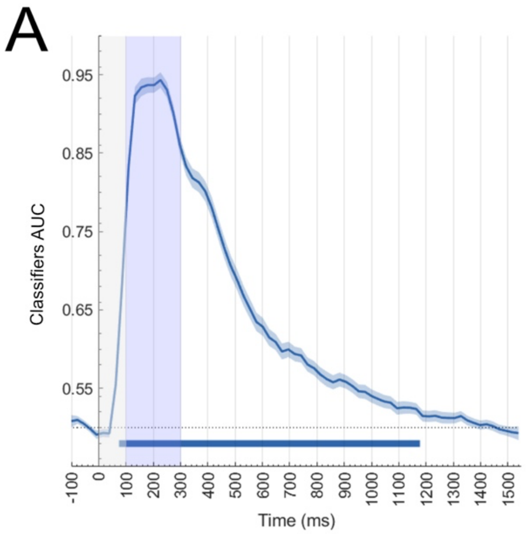


S1. **Canonical Template Tracking results.** Localizer's result after classifying faces and names. Grey shading indicates stimuli presentation. Blue shading shows the selected time window to create the two template patterns for faces and names.

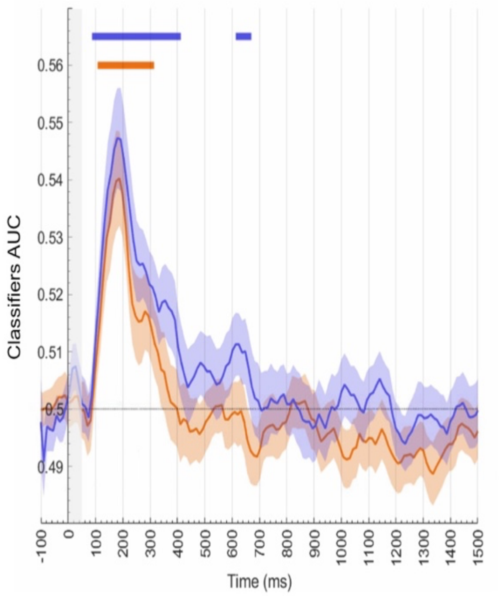


S2. **Raw voltage classification result using all cues to train and test the classifier.** Result of the time-resolved classification of the category (faces vs. names) to be selected (blue) vs. expected (orange). Horizontal colored lines indicate statistical significance against chance within each block. Grey shading indicates cue presence onscreen. Training a linear classifier with the two cues of each condition together to decode stimulus category separately for Attention and Expectation showed early peaks in classification that quickly vanished. We compared both AUC results by subtracting them and performing a one-side t-test against 0 (corrected for multiple comparisons with a cluster-based approach), which yielded no significant differences (all ps>0.05). To directly compare MVPA and RSA results, we performed a regression analysis using the RSA results as predictors of each participant's decoding accuracy. We then compared the variance explained by each regressor (i.e. RSA model) in a one-way ANOVA. Post-hoc comparisons revealed that, as expected, the Cue model explained the decoding results depicted in Figure S4 better than any other RSA model (all ps<0.001).

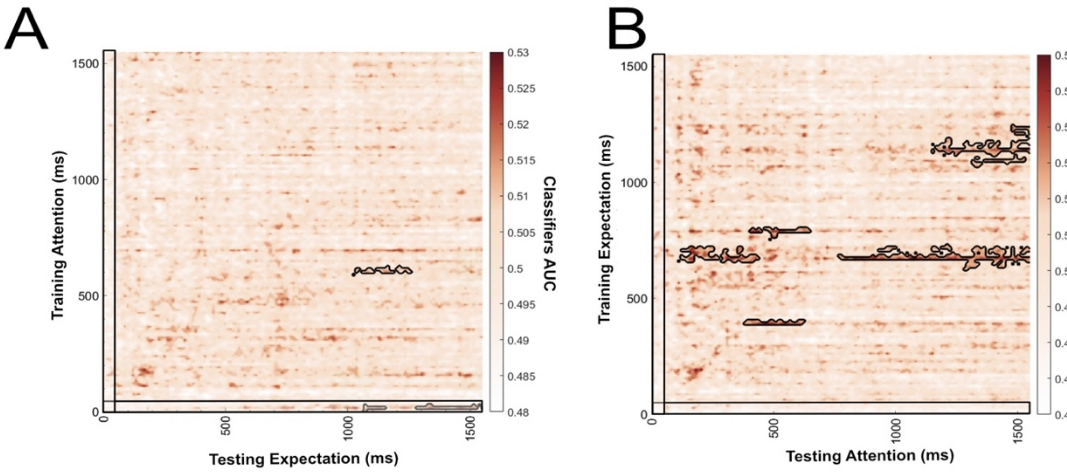


S3. **Cross-classification results on raw voltage split by training and testing direction.** (A) Train Attention-Test Expectation. (B) Train Expectation-Test Attention. Black outline depicts significant clusters.

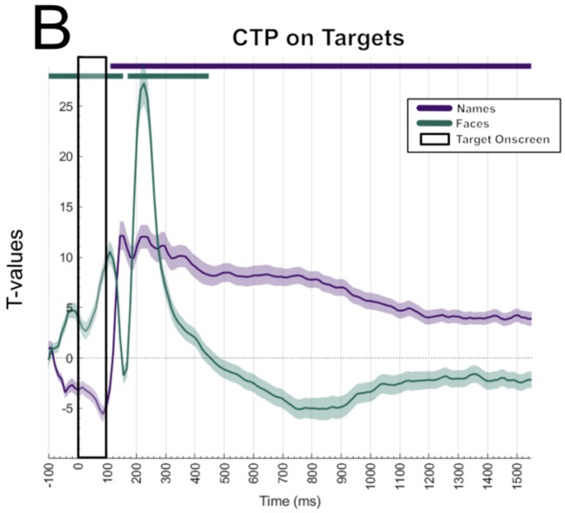


S4. **Canonical Template Tracking results.** (A) Localizer's result after classifying faces and names. Grey shading indicates stimuli presentation. Blue shading shows the selected time window to create the two template patterns for faces and names. (B) Canonical template pattern result for targets. Since there were no statistical differences between Attention and Expectation, the results are averaged. The green line shows the results for how much the face CTP explains face perception. Purple shows the result for the name CTP on name stimuli.

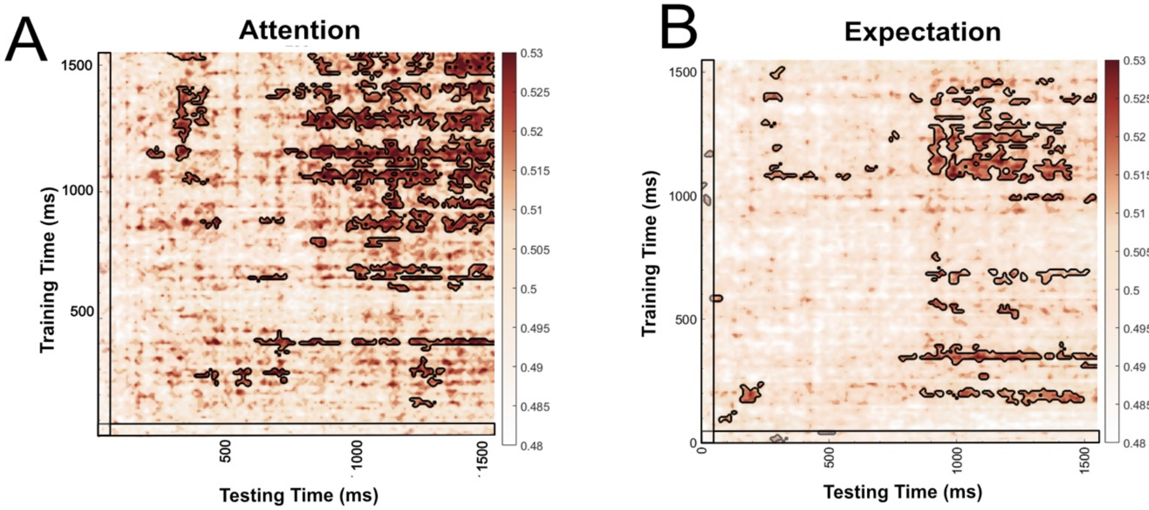


S5. **Classification results only using trials from different blocks** (A-B). Note that the patterns of generalization are stable in both conditions. However, the appear reduced, likely due the loss of power after using less trials to train and test.
